# FAMUS: A Few-Shot Learning Framework for Large-Scale Protein Annotation

**DOI:** 10.64898/2026.03.08.710366

**Authors:** Guy Shur, David Burstein

## Abstract

Predicting gene function is a pivotal and challenging step in genomic and metagenomic data analysis. Current automatic annotation tools typically rely on the single most similar sequence from the query database and struggle to robustly set hit thresholds for annotation. The sparsity of proteins per annotation makes it challenging to confidently assign gene function for underrepresented families. Here, we present a contrastive learning framework for functional annotation. FAMUS (Functional Annotation Method Using Supervised contrastive learning) compares query sequences to a full array of profile Hidden Markov Models and transforms the similarity scores into a condensed vector space that minimizes the distance of proteins from the same family. The similarity scores of a query to all profiles are used for its representation instead of considering only the top-ranking hit. Unannotated sequences are incorporated as negative examples during training, enabling robust detection of proteins that fall outside the scope of the reference database without requiring a user-defined threshold. Using this approach, FAMUS outperformed KEGG’s native KofamScan for KEGG Orthology annotation and InterPro’s InterProScan for PANTHER family annotation. We thus created four protein annotation models using protein families from the KEGG Orthology, InterPro family, OrthoDB, and EggNOG databases. All four models are available as a conda package and via our user-friendly web server, allowing users to annotate large-scale datasets. FAMUS is the first comprehensive and modular annotation framework based on contrastive learning. It supports both pre-defined and user-specific databases for tailored annotation, and can be easily integrated into any genomic and metagenomic analysis pipeline to facilitate accurate, large-scale functional annotation.

## Introduction

Advances in sequencing technologies and bioinformatic approaches over the past decades have greatly increased the rate at which we can sequence and assemble genomes and metagenomes. A key informative analysis following the assembly is gene functional annotation, where each putative gene is assigned a function based on an annotated database. The most widely used tools for gene annotation are based on sequence similarity, from simple BLAST^1^ searches to deep learning and state-of-the-art language models^2–6^. One of the sensitive and efficient approaches to functional annotation involves using profile Hidden Markov Models (pHMMs), statistical models that capture position-specific patterns in sequence families. Since these models represent families composed of multiple sequences, they can capture complex and subtle sequence patterns, eliminate the need for pairwise sequence alignment, and reduce the total number of comparisons required to make a prediction compared to sequence-sequence search approaches^7,8^. Gene sequences are compared to pHMMs to acquire measures of similarity to the protein family represented by the profile.

Although straightforward and effective, three major caveats to this approach limit its accuracy. First, annotation is typically based on the best score (“winner takes it all”), and this strategy does not make full use of all the bit score information generated in the search process against all the families in the database. In most cases, using the highest similarity scores is adequate, but accounting for entire similarity patterns may allow better annotation of distant homologs and ambiguous cases. Second, for both orthology- and function-based pHMMs, performance is affected by the criteria used to cluster proteins into families. E.g., pHMMs are prone to the inclusion of genes with low intra-cluster similarity. This could lead to low-information pHMMs with low specificity, yielding high rates of false positives or low rates of high-scoring hits. Third, defining a reliable annotation threshold is non-trivial, as optimal cutoffs vary across protein families due to differences in evolutionary rates, sequence diversity, and family size, often requiring manual curation or family-specific tuning.

These challenges are well exemplified by the Kyoto Encyclopedia of Genes and Genomes (KEGG)^9–11^, a comprehensive knowledge base for systematic analysis of gene functions, linking genetic data with higher-order functional information through standardized pathway maps and molecular networks. KEGG Orthology is a database of functional gene orthologs, also termed KOs. In KEGG, human curation is a necessary part of the KO assignment process, since this assignment is often context-dependent, and expert biological insight might contradict the relationships learned by automated annotation tools. The result is a significant number of KOs with highly diverse sequences in terms of similarity, which are difficult for automation tools to generalize. For example, despite their distinct functions, the KO labeled K13194 includes sequences of both ADARB1, a double-stranded RNA editing enzyme, and ADARB2, a catalytically inactive homolog that is believed to be a competitive inhibitor of ADARB1^12^. Such cases are expected to lead to pHMMs with low specificity, as described above. Applying more advanced classification methodologies is also problematic, as KEGG contains more than 24,000 families, making the training of accurate multi-class classifiers challenging. Classifying sequences to a KEGG protein family directly would require either a very complex model or individual models for each family. Both approaches require substantial computational resources, and with 20% of all KOs having 100 or fewer sequences, there are not enough positive examples for many KOs for direct classification. The KEGG-native annotation tool, KofamKoala, addresses this using pHMMs with adaptive score thresholds, yet remains subject to the caveats described above. These challenges are not unique to KEGG; analogous issues affect other databases such as InterPro^13^, EggNOG^14^, and OrthoDB^15^, all of which contain families with highly variable sequence diversity and sparse representation.

Similar challenges were encountered in other fields, such as computer vision, where a common task is to annotate facial images with the identity of the person given a single or very few images. Contrastive learning has emerged as an elegant solution to facial image annotation, comprising methods such as Siamese Neural Networks (SNNs)^16^ and later triplet networks^17^. Contrastive learning models can learn effectively even with very limited training data by transforming the task of classification to a task of comparison. In the training process, a single neural network is used to generate a low-dimensional representation of different objects and is optimized such that objects with the same label are co-located in the embedding space. This dramatically increases the number of possible samples for training due to the combinatorial nature of sampling multiple objects and decreases the dimensionality of the output space, alleviating the need for extensive models. Lifted Structured Feature Embedding^18^ extended this paradigm by leveraging all positive and negative pairs within a batch rather than individual triplets, yielding a more stable and informative training signal. Later methods, such as Supervised Contrastive learning (SupCon)^19^ replaced the margin-based loss with a temperature-scaled, normalized softmax, giving finer control over how much each pair contributes according to its difficulty. Previous approaches that utilised contrastive learning for biological sequence analysis include SENSE^20^, a tool that encodes *k*-mer presence and position for alignment-free sequence embedding and pairwise distance estimation; CLEAN^6^, a tool that encodes transformer model outputs for enzyme classification using both SupCon and triplet networks; and ATGO^3^, a triplet-network-based model for Gene Ontology (GO) annotation. CLEAN and ATGO both use representations derived from language models as their contrastive model’s input. While useful, the preprocessing step of using language models in CLEAN and ATGO represents a significant computational overhead that limits the scale at which they can be used to annotate a large number of proteins. In addition, they were both developed and evaluated on one type of annotation (EC numbers^21^ and Gene Ontology (GO)^22^, respectively), and were not designed to support user-defined custom databases or to identify sequences that fall outside the scope of the reference database. Thus, a general and scalable approach is needed to advance our understanding of environmental microbes’ functional potential.

We adopted a SupCon-based approach to address the challenges of full genome/metagenome functional annotation, which we call “Functional Annotation Method Using Supervised contrastive learning” (FAMUS). FAMUS explicitly classifies sequences to a specific protein family using a neural network trained with SupCon, or identifies them as out-of-scope when they are not sufficiently similar to any family in the reference database. This is achieved by incorporating unannotated protein sequences as negative examples during training, enabling robust out-of-distribution (OOD) detection without requiring a user-defined threshold. Since many protein families represent only a small fraction of the total annotated sequences, the few-shot learning capability of the SupCon-based model is a key component of our algorithm and produces highly accurate assignments. FAMUS is modular by design, supporting both pre-built models for four major databases and user-defined custom databases, making it applicable to diverse biological domains. We validated and benchmarked our approach using the KEGG^11^ Orthology database, PANTHER^23,24^ protein families, and traditional pHMM-based annotation. We further provide both a conda package and a web server that implements our approach to assign proteins to KEGG Orthology, as well as InterPro^13^ protein families, OrthoDB^15^ groups, and a combined dataset of COGs^25^, KOGs, and arCOGs^26^ from EggNOG^14^.

## Methods

### Data acquisition

All KEGG^9–11^ proteins, along with the orthology database, were downloaded from the KEGG FTP server^27^. KEGG data used to train the model for evaluation was downloaded on May 14^th^, 2021, and KEGG data used to test the model and to train the latest version of the model was downloaded on May 24^th^, 2023. PANTHER versions 18 and 19 sequences and annotations were downloaded from the PANTHER database website. UniRef90^28^ protein sequences and ID mappings used to generate the InterPro, EggNOG, and OrthoDB models were downloaded on January 14^th^, 2024 from the UniRef FTP server^29^.

### Generating unlabeled sequence sets

FAMUS uses unlabeled sequences for training and inference to label sequences as either one of the previously seen protein families or as “unknown”. For KEGG, these sequences were taken from the KEGG Genes database if they were not assigned to any KEGG Orthology label. For PANTHER, UniProt KnowledgeBase IDs of unannotated sequences from the organisms annotated by PANTHER were gathered (according to https://pantherdb.org for PANTHER 19 and https://archive.pantherdb.org for PANTHER 18). Each unannotated sequence’s UniProt KnowledgeBase ID was mapped to a UniRef90 sequence. For InterPro, OrthoDB, and the EggNOG models, these were UniRef90 sequences whose UniProt ID was not mapped to any InterPro family, OrthoDB family, and any COG, KOG, or arCOG family, respectively.

### Generating sub-families and pHMMs

We created two model versions for each database: a comprehensive version with ortholog families clustered into sub-families and a “light” version where only exceptionally large ortholog families were clustered into sub-families (see below), resulting in substantially smaller pHMM databases. To create sub-families, we sub-clustered orthologous groups using mmseqs2^30^ cluster (version bad16c765aac60d84a8fde3548adbb06b34980bd). Each set of sequences was clustered twice, as detailed below. The first clustering aimed to reduce sequence redundancies within each protein family. The non-redundant sequences were clustered again to partition the protein families into sub-families.

The pipeline to partition families into sub-families is as follows. For each family:

1. If the family contains fewer than five sequences, skip sub-clustering of this family.
2. Remove redundant sequences: Cluster all the protein sequences using mmseqs2 cluster with parameters sensitivity=7.5, coverage=0.8, and sequence-identity=0.9. If the data are being processed for the light version of the model, and this is not an extensive family (specifically, the length of the longest representative sequence times the number of representative sequences is less than 500,000,000), skip the next steps and use all representative sequences as a single cluster representing the family.
3. Cluster the representative sequences to sub-families using mmseqs2 cluster with parameters sensitivity=7.5 and coverage=0.5.
4. Sort the resulting clusters by the number of sequences. Starting with the largest cluster in descending order, create a sub-family from each cluster until the cumulative number of sequences in the sub-families covers at least 80% of the representative sequences.
5. Generate pHMMs for each sub-family that passed the previous step. If any clusters remain after covering 80% of the representatives (“leftover” sequences), search these sequences against the existing pHMMs and add each sequence to the sub-family of the pHMM with the highest bit score.
6. If any “leftover” sequences did not match any profile (no search hits), create a single sub-family comprised of these sequences.

Sub-families with fewer than six sequences were artificially augmented by constructing a pHMM and using it to generate sequences from the profile using hmmemit (HMMER version 3.3.2) with default parameters. The purpose of this augmentation step is to increase the sample size of extremely small subclusters, allowing them to be split into three sets that are required for downstream steps. After the augmentation of small subclusters, pHMMs were constructed for each subcluster using hmmbuild. The number of sequences, families, and sub-families in each dataset is detailed in Supplementary Table S1.

In addition to KEGG and PANTHER pHMMs, we created pHMMs from UniRef90 sequences annotated with InterPro^13^ families, OrthoDB^15^ families with 200 or more sequences, and COGs, KOGs, and arCOGs families. To assign sequences to orthologous groups of these databases, we used mappings between UniRef90 and each database that were acquired from the UniRef FTP site^28^. All pHMMs were constructed by performing multiple alignments using MAFFT^31^ (version v7.475) with default parameters, followed by hmmbuild (HMMER version 3.3.2) with default parameters.

### Data sampling and pHMM search

To reduce potential bias induced by extremely large sub-families and decrease the hmmsearch runtime during model training, we created the training set for KEGG by randomly sampling up to 60 sequences from each family or sub-family (PANTHER families/sub-families were not sub-sampled due to the smaller size of this database). We further sampled unlabeled sequences from the non-redundant set to achieve equal numbers of labeled and unlabeled examples for both KEGG and PANTHER. The sub-family pHMMs were used to scan the training set using hmmsearch (HMMER^32^ version 3.3.2) default parameters. HMMER reports two types of bit scores: the full bit score, which is calculated by comparing the entire sequence to the pHMM, and the best domain bit score. Based on model performance, we chose to use the best domain bit scores as input. After running hmmsearch, the results were parsed to extract the best domain bit scores of each query against all the sub-families’ pHMMs. The result of this step is an *N*×*M* matrix, where *N* is the number of sampled sequences and *M* is the number of sub-families, which serves as the input for our model. To avoid data leakage, it was imperative to avoid using sequences that were part of the pHMM generation. To that end, we adopted the strategy used by KofamKoala: each sub-family was split into three equally sized groups (± one sequence). A pHMM was created based on two of the three subgroups, and this pHMM was used to score the sequences of the third subgroup. This process was repeated for every third of the sub-family to acquire an unbiased bit score, representing its resemblance to its own sub-family. Bit scores for sequence-profile pairs that were not reported in hmmsearch were regarded as zero.

### Model architecture

We trained a neural network (architecture depicted in Fig. 1) with a Supervised Contrastive (SupCon)^19^ loss function to create protein embeddings such that sequences of the same label were close to each other in the embedding space. Bit score vectors from the preprocessing phase are fed into the network to generate low-dimensional projections of the input data. The goal of SupCon is to optimize the parameters to minimize the distance between two instances with the same label and maximize the distance between instances with different labels. As a result, samples from the same class tend to be co-located within the embedding space. A critical challenge in protein function annotation is OOD detection, i.e., identifying when a protein does not belong to any family in the reference database. This is particularly important for frameworks that support custom models trained on arbitrary databases of a restricted functional domain. To improve OOD detection, we incorporated into the SupCon batches unlabeled protein sequences as negative examples alongside labeled training data.

**Figure 1.**
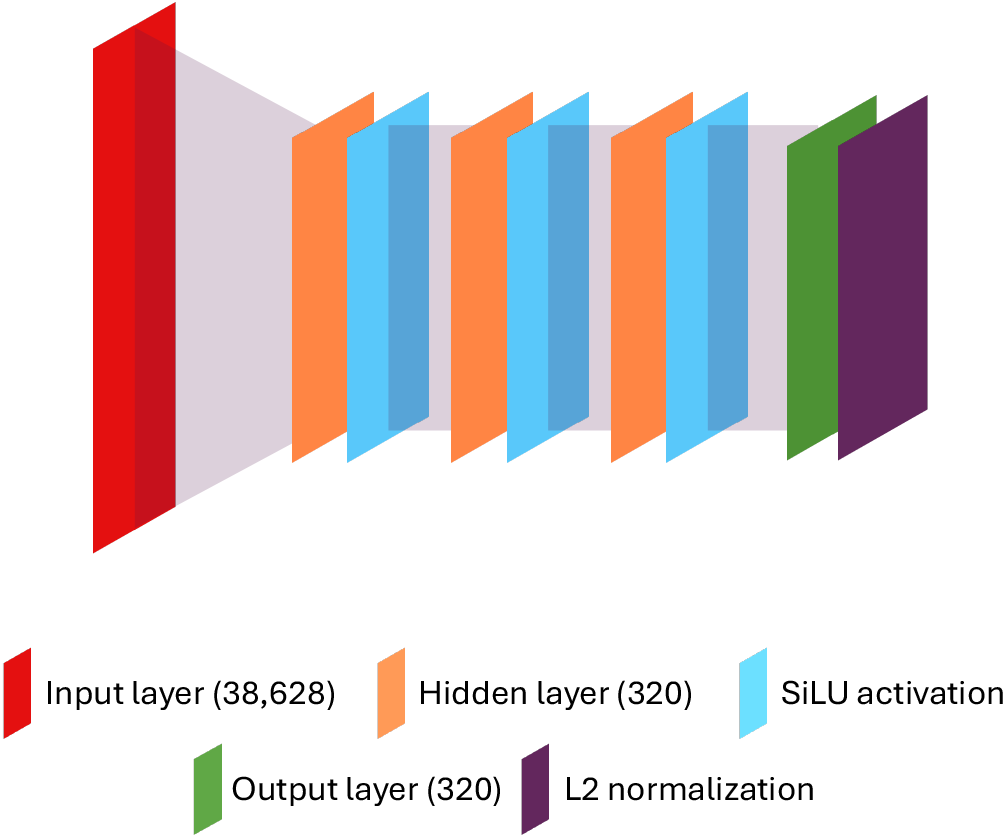
FAMUS neural network architecture. The size of the input layer corresponds to the number of sub-familiy pHMMs in the database (KEGG orthology in this example). Other layer sizes remain the same regardless of pHMM database.

All neural network models were trained using PyTorch^33^ version 2.2.0 and CUDA version 11.8. We used the sum-inside-the-log SupCon loss function to train the model. Briefly, the augmentation-free SupCon loss function for a single anchor (labeled sample) *i* used in FAMUS is defined as follows:

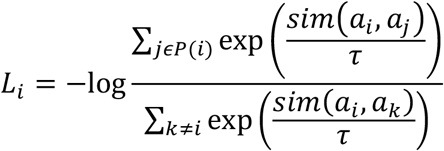

Where *a*_*i*_ is an embedding generated by the network for sample *i, sim* is the similarity function used to evaluate projection distance (in our case, *sim*(*x, y*)= *x* · *y*, the dot product of *x* and *y*), *P*(*i*)is the set of all samples with the same labels as in the batch excluding *i* itself, and τ (temperature) is a positive real number and a hyperparameter of the model. Since our network normalizes the projected vectors to length 1, the similarity of the raw projections used to train the network is equivalent to cosine similarity: the Euclidean norm of *x* ™ *y*. The total loss of one batch *L*_*b*_is thus:

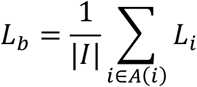

Where *A*(*i*)are the labeled samples in the batch and |*I*| is the number of valid anchors (with ≥ 1 same-label samples) in the batch,.

We used the PyTorch Adam optimizer with default parameters, no weight decay, no drop out, and a learning rate of 10^−5^, which decreases by 50% every 15 epochs.

Our model consists of an input layer of size *M*, equal to the number of sub-families, three hidden layers of size 320, and an output layer of size 320. Batches were created by randomly sampling 16 labels, and up to eight samples were drawn from each of these labels. In addition, 64 unlabeled samples were added to each batch for a total batch size of 96–192 samples. The sizes of the hidden and output layers and the number of layers were chosen based on classification performance after a grid search on training data split to training and evaluation sets over layer sizes 250, 300, 310, 320, 330, 340, and 350, one to four hidden layers. In addition, the embeddings are normalized to a hypersphere using an L2 normalization layer. We apply the SiLU^34^ activation function to the output of the hidden layers.

### Evaluation metrics

The F1 score, or F-measure, was calculated as:

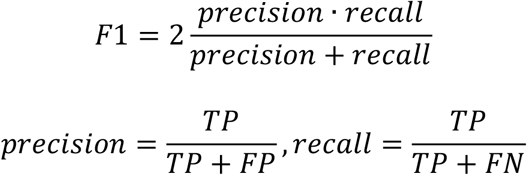

Where *TP, FP*, and *FN* are the number of true positives, false positives, and false negatives, respectively. The micro F1 score was calculated as above, without considering class size imbalances. The weighted F1 score is calculated by taking the average of F1 scores over each class, weighted by the number of sequences in each class.

### Sample inference and performance evaluation

To infer labels of input sequences after training, it was necessary to contrast their model-generated embeddings to those of functionally annotated sequences. For this purpose, all sequences used for training were transformed to embeddings using the model after training was completed. Input sequences for annotation were transformed to bit score vectors with the same pHMMs used for training. These vectors were then encoded using the trained model, and the Euclidean distance between the input embedding and that of every training sequence was calculated. If the nearest neighbor is assigned to a family, and its distance from the input sequence encoding is less than a pre-calculated threshold, the input sequence is assigned to the same family (or families, if the nearest neighbor belongs to more than one family). Otherwise, the sequence was labeled as “unknown”. The global threshold is calculated prior to classification using the trained network in three-fold cross-validation: The training data is split into three, and for each fold, one of the three splits serves as the validation data, and the other two make up the new training data. The shortest distance between each training sample and any validation sample is calculated, and the selected threshold for the validation set is the one that maximizes the weighted F1 score in the classification of the validation dataset. The final global threshold used was the average of the three selected thresholds. In the inference stage, FAMUS annotates sequences using the original protein family labels, and not sub-family membership.

To guarantee that no sequence appears in both the training and test sets of either KEGG or PANTHER, all labeled and unlabeled sequences in each training set were clustered using mmseqs linclust with 0.8 coverage and 90% sequence identity. The test sets were created to be the representative sequences from each cluster that did not contain any sequences existing in the older version of their database. The training and test sets for the OrthoDB database and the EggNOG database, comprised of the union of COG, KOG, and arCOG protein families, were created by first clustering each protein family using mmseqs linclust with 0.8 coverage and 90% sequence identity. For each set of representative sequences of each family with at least two representative sequences, 20% (rounded up) were set aside for the test sets.

KofamScan^7^ and InterProScan^35^ do not explicitly annotate sequences; rather, they provide a list of possible annotations. To calculate the F-score for KofamScan, we set its prediction for each sequence to be the one with the highest bit score that has passed the KofamScan pHMM significance threshold, if one exists, and “unknown” otherwise. For InterProScan, we set its prediction for each sequence to be the one with the lowest E-value in the output if one exists, and “unknown” otherwise. KofamScan was used with profiles from June 1^st^, 2021, downloaded from the FTP server^27^. InterProScan version 5.66-98.0 was downloaded from the FTP server^36^ and used with every available analysis, residue level annotation disabled, only PANTHER database search enabled, and without identical sequence lookup.

### Runtime evaluation

All runtime tests were performed on the same hardware with 24 virtual CPU cores on a Rocky Linux release 9.5 (Blue Onyx) machine with an AMD EPYC 7443 24-core Processor and an RTX A6000 GPU (10,752 CUDA cores).

## Results

FAMUS addresses three key challenges of pHMM-based annotation: (1) it leverages all similarity scores against the full profile array rather than relying solely on the best hit; (2) it accurately annotates even sparsely represented families through few-shot contrastive learning; and (3) it identifies out-of-scope sequences without requiring a user-defined threshold, by incorporating unannotated sequences as negative examples during training. As an initial test case, we applied FAMUS to KEGG Orthology annotation (Fig. 2).

**Figure 2.**
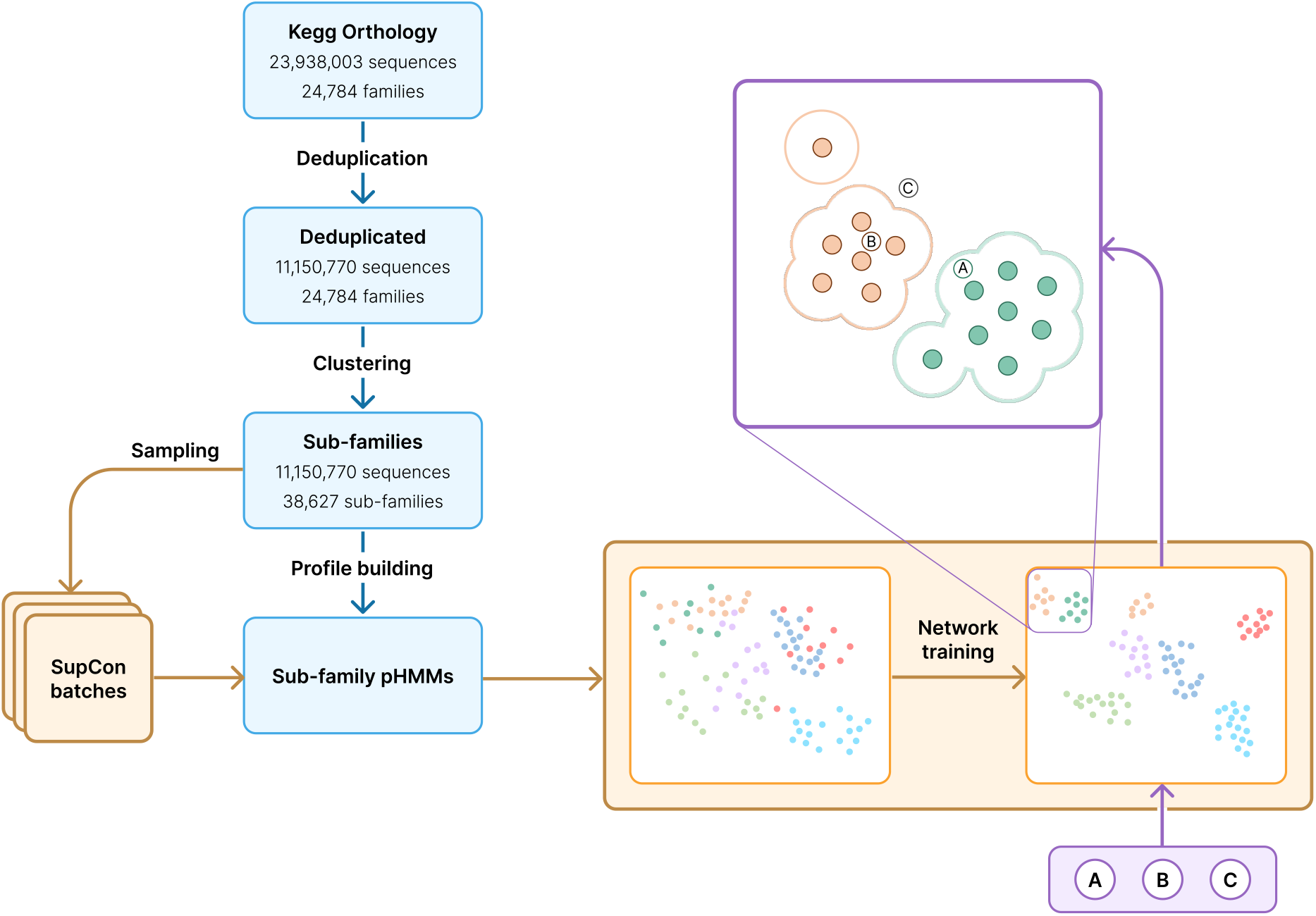
The FAMUS pipeline for end-to-end model training and sample classification exemplified on KEGG Orthology. Each protein family is automatically filtered for redundancies and clustered into sub-families that are used to generate pHMMs. Sequences sampled from the training set are scored against each pHMM, and the score vectors are used in batches to train the neural network for embedding generation. For inference, the model assigns the label(s) of the nearest training set embedding if sufficiently close in the embedding space.

### Data preprocessing

As a preprocessing phase, highly similar sequences within each protein family were filtered out to remove redundancies due to representation biases in the database (Fig. 3). Next, each non-redundant sequence was sub-clustered to generate high-resolution protein sub-families (Fig. 4), and a pHMM was created for each sub-family. The gene embedding for the training set was generated by sampling sequences from each sub-family and scoring them against all the pHMMs. The vector of bit score per pHMM was used as the numeric representation for the neural network’s input.

**Figure 3.**
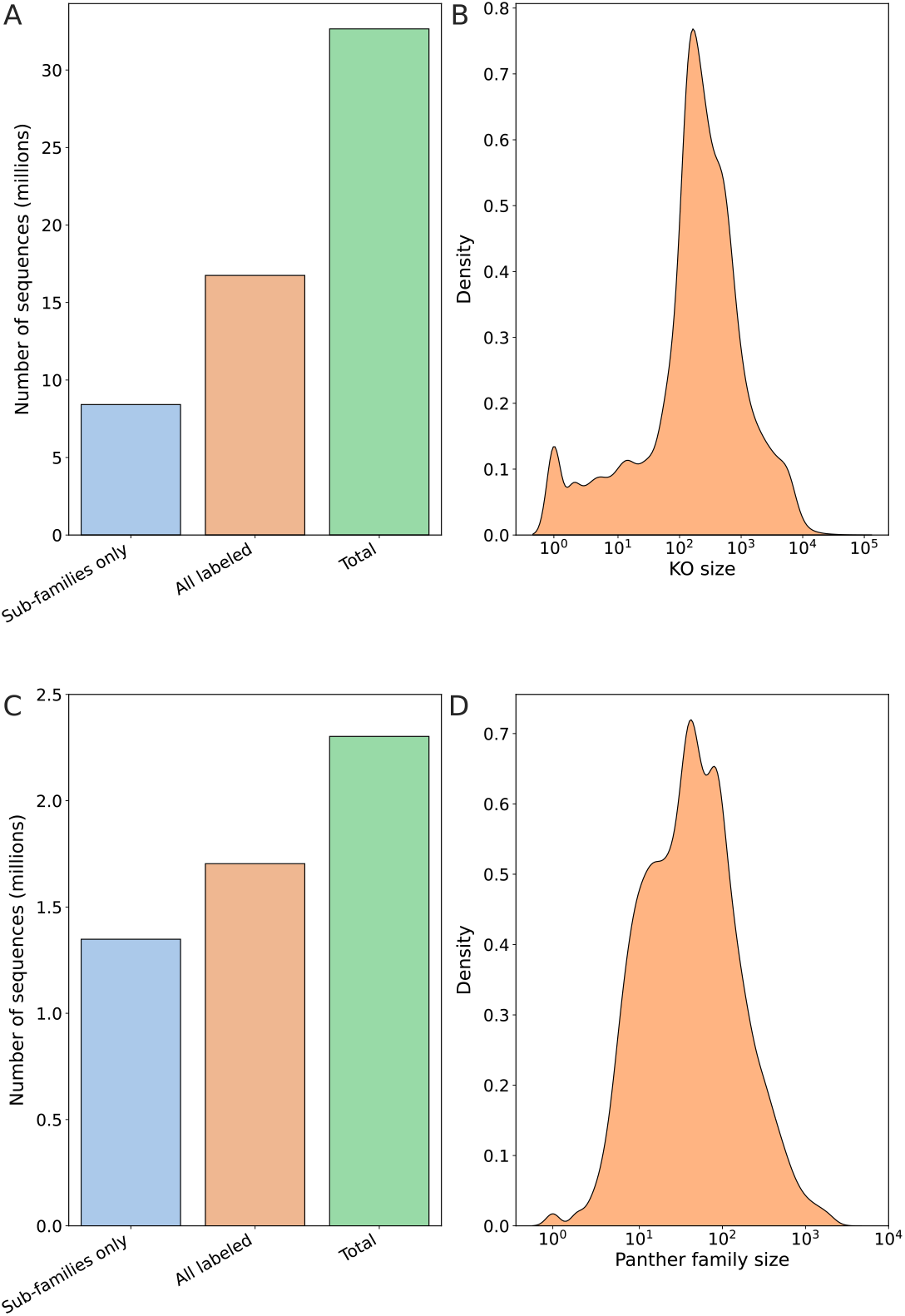
Distribution of protein sequences in KEGG Orthology (KO) and PANTHER protein families. A. The number of sequences (in millions) that were selected to compose the sub-families after de-duplication, the total number of sequences that have a KO identifier before de-duplication, and the total number of protein sequences in KEGG (with or without a KO identifier) as of May 24^th^, 2023. **B**. Distribution of the number of proteins per KO (before de-duplication) on a logarithmic scale. **C**. The number of sequences (in millions) that were selected to compose the sub-families after de-duplication, the total number of PANTHER v18 sequences before de-duplication, and the total number of protein sequences in PANTHER v18 **D**. The distribution of the number of proteins per PANTHER family (before de-duplication) in logarithmic scale.

**Figure 4.**
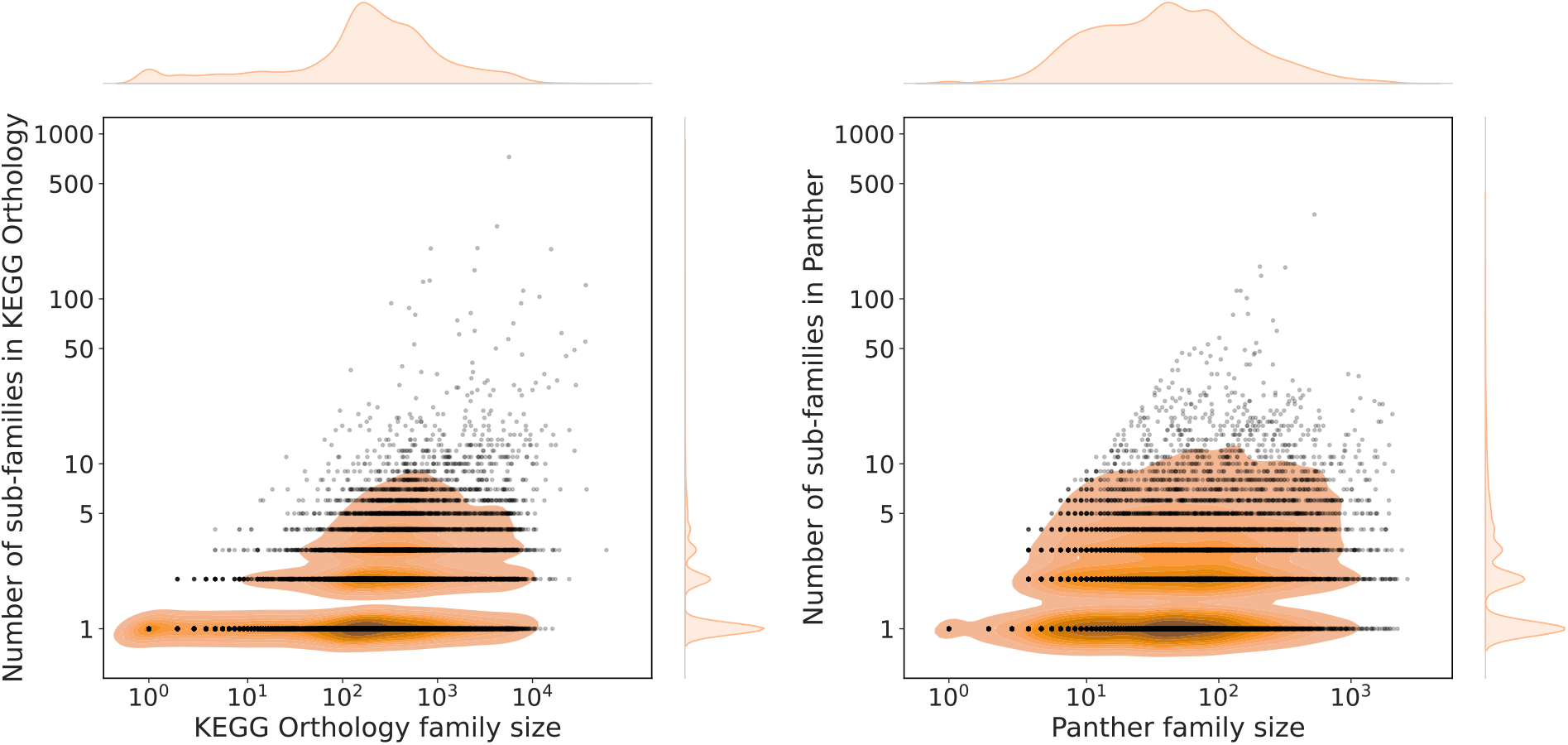
Number of sub-families vs. family size. Distribution of the number of sub-families in a KO family and PANTHER family with respect to the family size (i.e., number of proteins per family) before de-duplication. Each sub-family is represented by one dot. Both axes are on a logarithmic scale.

To prevent data leakage, sequences should not be scored using profiles derived from them. To that end, we adopted the strategy used by KofamKoala^7^: each sub-family was split into three equally sized groups (±1 sequence). A pHMM was created based on two of the three subgroups, and was used to score the sequences in the third subgroup (which was not used to build the pHMM). This process was repeated three times to acquire an unbiased bit score representing the sequence’s resemblance to its own sub-family.

### Classification

To label new input sequences, first, their similarity scores to the full set of sub-family profiles were acquired. Using these scores, the neural network generated embeddings in a lower-dimensional space. For the KEGG model, this meant reducing the input from a vector of 38,628 bit-scores, one for each KO sub-family pHMM, to a 320-dimensional space. During the inference phase, the network was used to generate embeddings for individual sequences. The label prediction for a sequence was based on its nearest neighbor in the embedding space. Sequences were classified as ‘unknown’ if their embeddings were closest to an unlabeled training sample or if their nearest neighbor was further away than an empirically precalculated global threshold (see Methods).

Since the pHMM search phase was the primary computational bottleneck for the classification times, we created a light version of the model using fewer pHMMs to improve computational efficiency for large-scale analyses. In this light version, each protein family (KO in the case of KEGG, PANTHER family in the case of PANTHER) was represented by a single pHMM, instead of being clustered into multiple sub-families. This lower-resolution approach significantly reduced computation time, while maintaining robust accuracy, making it particularly suitable for analyzing very large datasets.

### Classification evaluation

We evaluated our framework using two independent models trained on separate datasets. The first was trained on a KEGG Orthology database snapshot from May 2021, and the test set comprised 8.2 million de-duplicated protein sequences added to the KEGG database between May 2021 and May 2023, out of which 3.49 million proteins were labeled with a KO, and the rest 4.71 million genes were unlabeled (had no KO associated with them). The second model was trained using the PANTHER protein family database. The unlabeled sequences in both datasets were used as out-of-distribution representatives. We selected KEGG and PANTHER based on three criteria: sparse multi-labeling; availability of explicitly unannotated sequences; and the persistence of older versions of both databases and annotation tools (KofamScan and InterProScan), enabling reliable evaluation based on annotation of sequences by models and trained on older versions. We also evaluated FAMUS models trained on OrthoDB protein families and a unified set of COG, KOG, and arCOG families, though these two databases were deduplicated and split into train and test sets in a standard, non-temporal approach (Supplementary Fig. S2, Supplementary Tables S4-S5).

To evaluate the classification performance in different scenarios, we randomly sampled five sets of 5,000 sequences from the test sets with varying fractions (5% – 95%) of unlabeled sequences (Fig. 5). Each sample was classified using the comprehensive model, the light model version, a baseline classifier using best pHMM hits (Supplementary note 1), and KofamScan^7^ for KEGG or InterProScan^35^ for PANTHER. We calculated the average weighted F1 scores over the five samples for each fraction of unknown sequences (Fig. 5, Supplementary Tables S2–S3). Both the comprehensive and light versions of FAMUS consistently performed as well or better than KofamScan. InterProScan achieved higher F1 metrics when unlabeled sequences composed 5% or 25% of the test set, while FAMUS achieved higher metrics when they composed 50%, 75% or 95% of the dataset (which is more representative of real-world genomic and metagenomic data). The comprehensive (high-resolution) version of FAMUS, using sub-families, either outperformed or performed similarly to the light (low-resolution) version of FAMUS without sub-clustering families in all cases. In every scenario, excluding the highest fraction of labelled sequences, FAMUS had similar or better performance than the baseline HMM classifier. Since the weighted F1 score is biased towards classes with smaller support, we additionally calculated the micro F1 score to understand how the fraction of unlabeled sequences directly affects the model’s accuracy (Supplementary Fig. S1, Supplementary Tables S2–S3). This analysis showed that FAMUS is less prone to mislabel sequences than it is to miss a correct label. To investigate how reliable each method is when it does explicitly annotate a sequence, we directly calculated their precision by treating correctly annotated sequences with ground truth labels as true positives and incorrectly annotated sequences as false positives, including sequences without ground truth labels (Supplementary Fig. S1, Supplementary Tables S2–S3). FAMUS had comparable precision to KofamScan and similar or slightly better precision compared to InterProScan in all scenarios. These results demonstrate that FAMUS is reliable when annotating a sequence at the cost of an increased false negative rate. This property is particularly desirable in metagenomic annotation, where large fractions of uncharacterized proteins increase the risk of spurious annotations in tools that lack a robust out-of-scope detection mechanism. However, when faced with a balanced mix of familiar and unknown function sequences, FAMUS achieves better performance than both Kofam and InterProScan.

**Figure 5.**
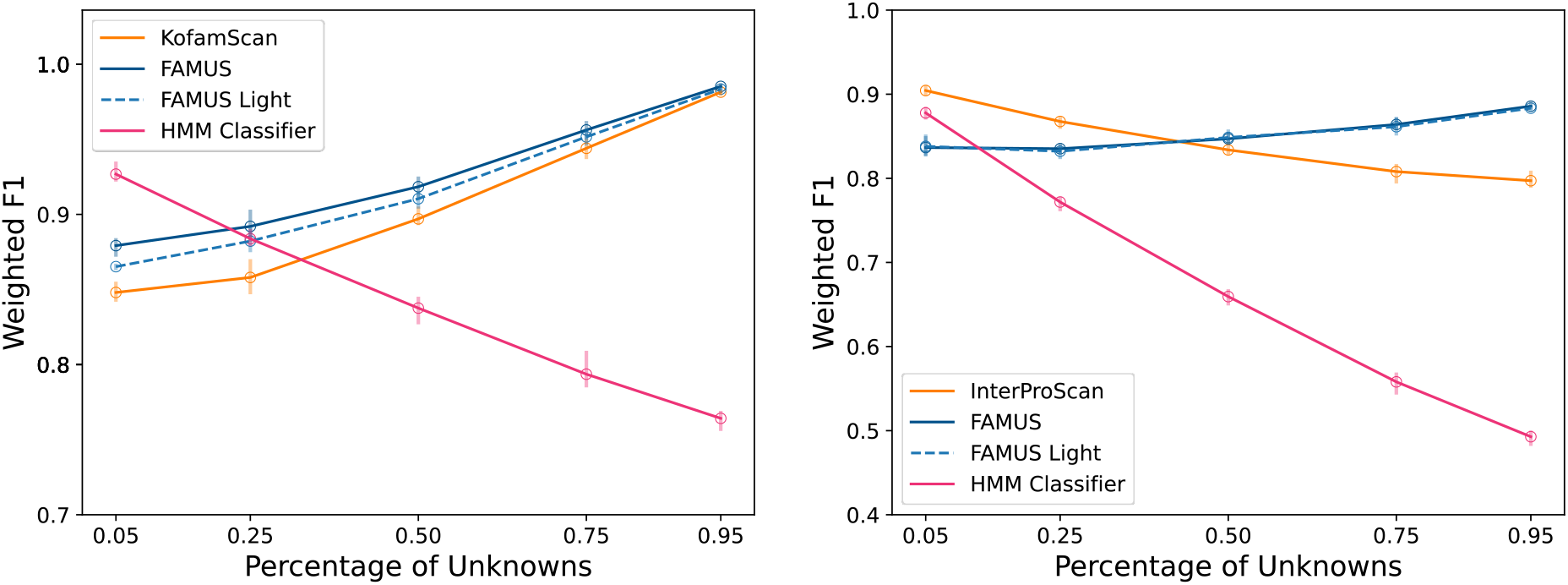
Classification performance of FAMUS compared to KofamScan and InterProScan: 5,000 sequences were randomly sampled five times from each test set using different fractions of unknown sequences (0.05, 0.25, 0.5, 0.75, and 0.95). Each sample was classified using the comprehensive and light implementations of FAMUS, as well as KofamScan for KEGG and InterProScan for PANTHER. The error bars represent the range of the five repeats of each labeled/unlabeled fraction.

### Runtime Evaluation

To compare the running time of each method, we sampled five sets of sequences of varying sizes (Fig. 6, Supplementary Tables S6–S7). The comprehensive and light models were tested on both CPU and GPU cores, as FAMUS was designed to support GPUs, while KofamScan and InterProScan were tested solely on CPU cores, since they currently do not support acceleration via GPUs. The runtime was bottlenecked by the pHMM search phase for both FAMUS and KofamScan. Due to the relatively small number of parameters in the model, using a GPU only yielded a marginal improvement in runtime performance.

**Figure 6.**
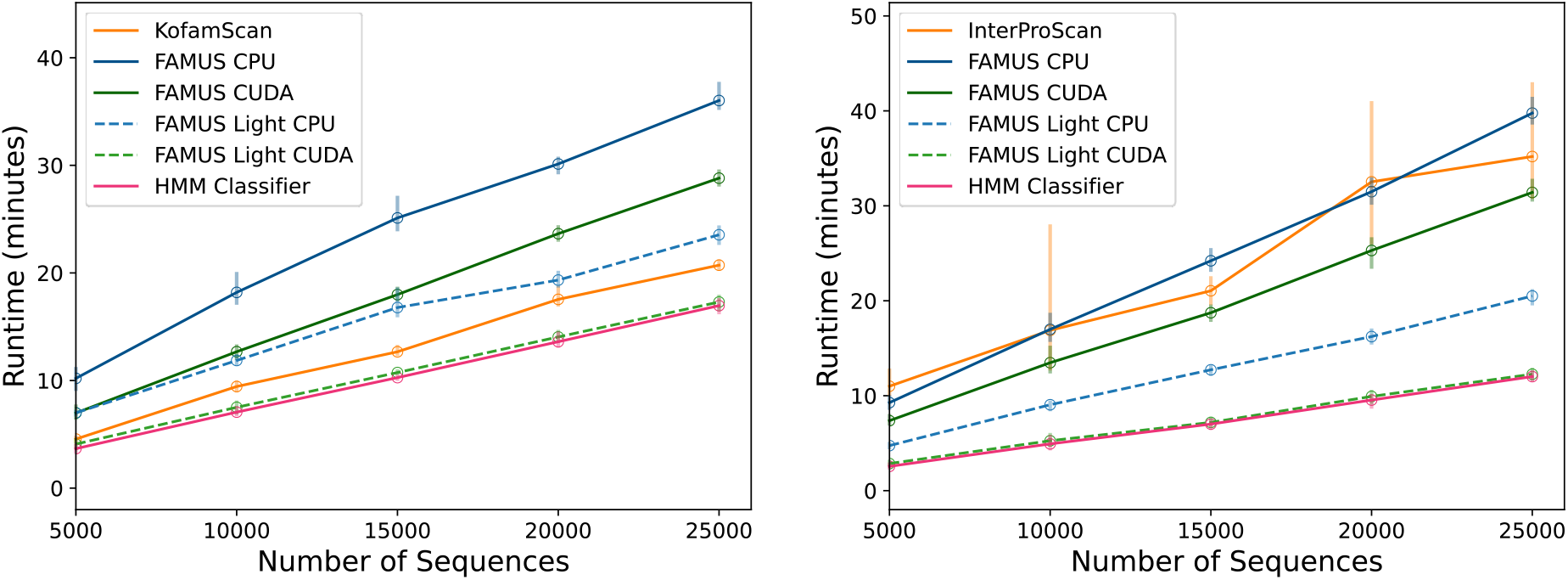
Runtime performance of FAMUS, KofamScan, and InterProScan. An increasing number of sequences were randomly sampled with ten repetitions from the test sets. Each sample was classified using the comprehensive and light implementations of FAMUS using KEGG and PANTHER with CPUs or a GPU, as well as KofamScan for KEGG and InterProScan for PANTHER. The error bars represent the range of the five repeats of each model.

FAMUS’s runtime is bottlenecked by the duration of sequence-to-pHMM comparisons, meaning the difference in runtime between the comprehensive and light models is proportional to the number of sub-families generated by the comprehensive model. In addition, the light model’s runtime is comparable to or better than any other pHMM-based annotation pipeline, as showcased in the comparison with KofamScan, InterProScan, and the baseline HMM classifier (Supplementary Tables S6–S7). The GPU-accelerated light model provides results faster than the benchmarked alternatives while also improving classification performance in most scenarios.

### Training additional FAMUS models

Following our approach’s performance, we used the same framework to train models for additional knowledge bases, namely InterPro families^13^, OrthoDB^15^, and EggNOG^14^. To provide easy access to our framework, we created a user-friendly web server that allows annotating FASTA files, available currently at https://app.famus.bursteinlab.org. These allow users to classify each of the input proteins based on combinations of the four trained databases using comprehensive or lightweight models: KEGG^9–11^ (38,627 comprehensive model families, 24,784 light model families), InterPro^13^ (102,590 comprehensive and 25,173 light model families), OrthoDB^15^ (67,414 comprehensive and 26,865 light model families), and EggNOG^14^ (70,417 comprehensive and 21,067 light model families). The results are presented in a tabular manner where sequences are assigned to a protein family out of each of the selected models, or annotated as “unknown” if classified as such by the model. Each column may be sorted and filtered to exclude certain results, and the (filtered or unfiltered) results may be downloaded directly. For large datasets and expert users, we open-sourced our code and provided all the pHMM databases compiled as part of this study. These high-resolution pHMM databases for KEGG, InterPro, OrthoDB, COG, KOG, and arCOG are publicly available at https://doi.org/10.5281/zenodo.14941373 and can be integrated into any HMM-based annotation pipeline independently of FAMUS.

Finally, we created a bioconda package named “famus” which provides the capability to download and install the aforementioned models locally, train custom models based on an arbitrary collection of protein families (with or without automatic protein family subclustering) and negative examples, and annotate protein sequences using any combination of pre-made or custom FAMUS models. Installation and usage instructions are provided in https://github.com/burstein-lab/famus.

## Discussion

We present FAMUS, Functional Annotation Method Using Supervised contrastive learning, which transforms protein functional annotation through a novel contrastive learning approach. By recasting the traditional annotation problem as a relational inference task, FAMUS overcomes fundamental challenges in genome-scale functional assignment, particularly the limited availability of training data for many protein families. Our framework begins with automated clustering of protein families to capture their inherent diversity, generating multiple profile Hidden Markov Models (pHMMs) that provide high-resolution similarity scores. These refined scores are then processed by a compact neural network, combining the sensitivity of pHMMs with the discriminative power of contrastive learning to achieve both high accuracy and sufficient speed for large-scale metagenomic analysis. Some databases, such as PANTHER^23^, do partition genes into sub-families, which are typically more homogeneous than the general families they comprise. Standardizing the use of a single computational tool (such as MMseqs2) to generate sub-families based on sequence similarity is an efficient and effective way to ensure that the pHMM landscapes derived from various databases lend themselves well to our framework, which emphasizes minimal user intervention. FAMUS directly addresses the three key limitations of pHMM-based annotation identified in the Introduction: best-hit reliance, low-specificity profiles for diverse families, and the difficulty of setting robust annotation thresholds at scale.

FAMUS was developed for general annotation, and it is thus applicable to any set of protein families and well-suited to annotating entire genomes and metagenomes. Our benchmarking results demonstrate that FAMUS is able to avoid misclassifying proteins while maintaining a high level of recall, a property that is particularly important when annotating metagenomes of understudied environments and species. Furthermore, FAMUS is modular by design, allowing seamless integration of various annotation databases and enabling users to leverage complementary information. This cross-database capability provides more comprehensive annotation coverage while maintaining consistency across different annotation sources. In addition, the streamlined and lightweight design of the FAMUS model allows for scalability, annotating millions of sequences in a short timeframe.

Like all machine learning approaches, FAMUS has inherent limitations in its ability to make novel functional predictions. While it can identify subtle and complex similarity patterns, the model still relies on its training data to make informative annotations. Additionally, practical constraints necessitated certain compromises in database coverage. For InterPro^13^, for example, our current model focused solely on families while excluding superfamilies to avoid an explosive growth in the number of sub-families. In the case of OrthoDB^15^, we included only groups with 200 or more members (26,865 families), and for EggNOG^14^, we restricted our scope to COG, KOG, and arCOG groups (21,067 families), excluding eNOGs. These choices were driven by balancing computational feasibility and output usefulness.

Future improvements to FAMUS could include a ranking system and confidence levels for predicted annotations. Such a system would let users filter predictions based on their specific confidence requirements, enabling flexible tradeoffs between annotation coverage and reliability. Owing to the modularity of our framework, users can produce different databases in various scopes, granularities, and for different domains. Additionally, with sufficient computational resources, exploring FAMUS’s performance on complete ortholog databases, including domains and uncharacterized proteins, could reveal the full potential of our approach for comprehensive annotation and organization of protein families.

## Data availability

The profiles (pHMMs) of KEGG, InterPro, EggNOG, and OrthoDB generated in this study, as well as the raw bit scores of sequences used to train the models, are available via Zenodo at https://zenodo.org/records/14941373. The web server developed as part of this study is available at https://app.famus.bursteinlab.org.

## Code availability

All code used to generate the models and this study, and that may be used to generate new models, is available via GitHub at https://github.com/burstein-lab/famus/. Associated pHMMs, model parameters, and other data required to use the pre-trained models for inference are available via Zenodo at https://zenodo.org/records/14941373 with installation instructions provided in the GitHub repository.

## CRediT author statement

**Guy Shur:** Conceptualization, Formal Analysis, Investigation, Software, Visualization, Methodology, Writing – Original Draft, Writing – Review and Editing. **David Burstein:** Conceptualization, Software, Methodology, Writing – Original Draft, Writing – Review and Editing, Supervision, Funding Acquisition.

## Acknowledgements

This study was supported in part by the Israel Science Foundation (grant number 355/23) and a fellowship from the Edmond J. Safra Center for Bioinformatics at Tel-Aviv University. We wish to thank Dr. Danielle Miller and Ella Rannon for their constructive feedback on this study, and Iris Burstein for help with the illustrations.

