## Supplemental Materials for "FAMUS: A Few-Shot Learning Framework for Large-Scale Protein Annotation"

*Supplementary data*

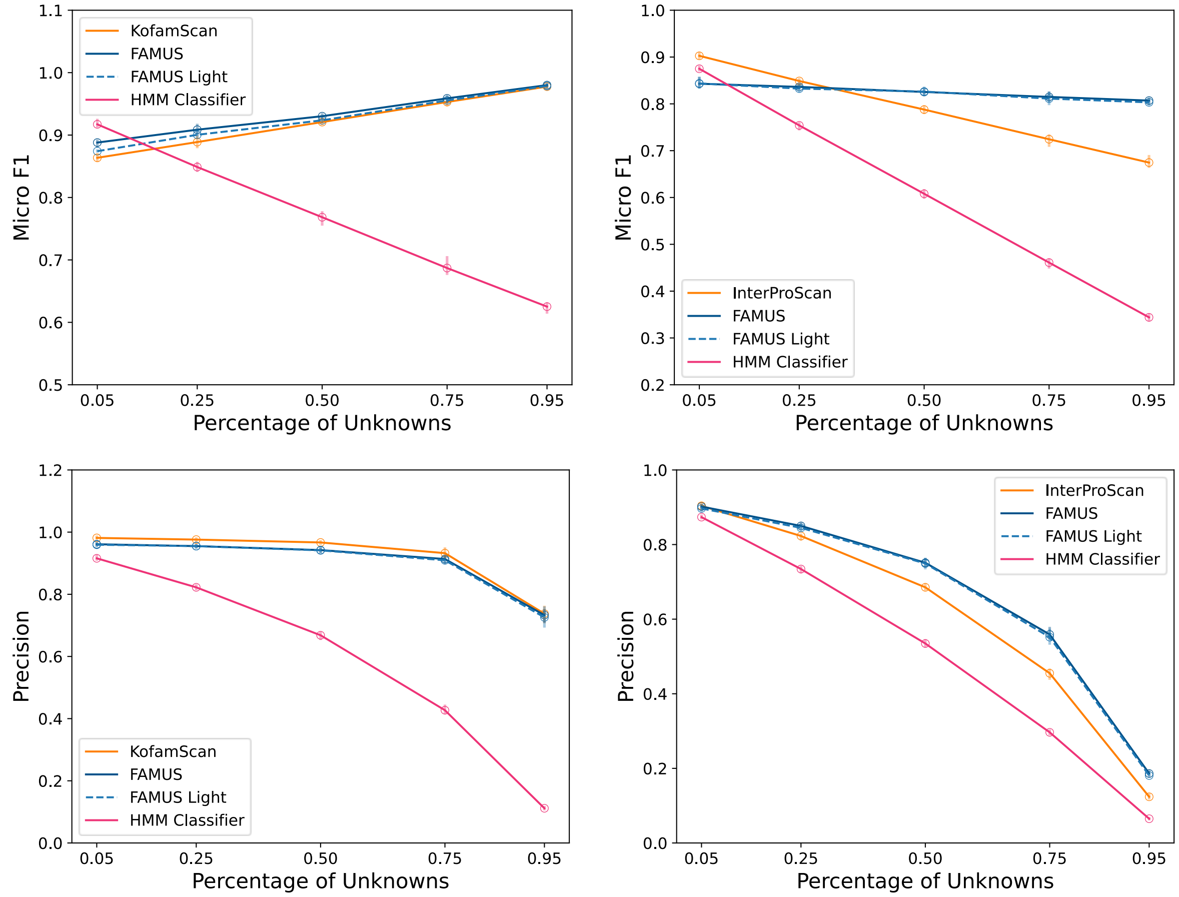


**Figure S1 | Micro F1 Classification performance and precision of FAMUS compared to KofamScan and InterProScan**: 5,000 sequences were randomly sampled five times from each test set using different fractions of unknown sequences (0.05, 0.25, 0.5, 0.75, and 0.95). Each sample was classified using the comprehensive and light implementations of FAMUS, as well as KofamScan for KEGG and InterProScan for PANTHER. The error bars represent the range of the five repeats of each labeled/unlabeled fraction.

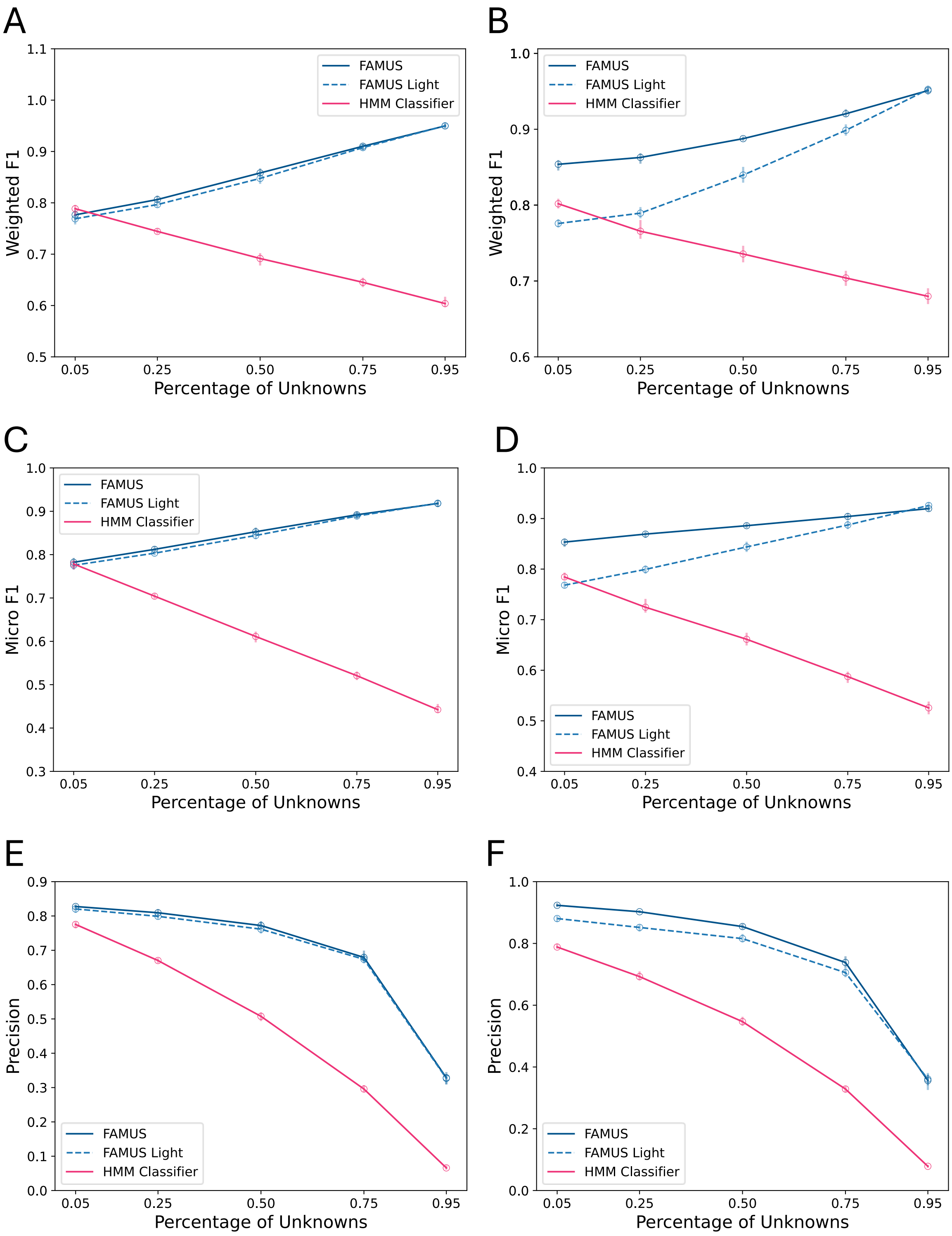


**Figure S2 | Weighted F1, Micro F1, and precision of FAMUS-OrthDB and FAMUS-EggNOG**: 5,000 sequences were randomly sampled five times from each test set using different fractions of unknown sequences (0.05, 0.25, 0.5, 0.75, and 0.95). Each sample was classified using the comprehensive and light implementations of FAMUS. The error bars represent the range of the five repeats of each labeled/unlabeled fraction. **A)** OrthDB weighted F1. **B)** EggNog weighted F1. **C**) OrthoDB micro F1. **D)** EggNOG micro F1. **E)** OrthoDB precision. **F)** EggNOG precision.

**Table S1:** Number of sequences before and after deduplication, and total protein families before and after clustering in each model.

| Database | Labeled sequences | Unlabeled sequences | Representative sequences (comprehensive) | Representative sequences (light) | Total sub-families (comprehensive) | Total families (light) |
|---|---|---|---|---|---|---|
| **KEGG (May 2021)** | 16,750,741 | 15,919,153 | 8,380,564 | 8,416,391 | 37,402 | 23,967 |
| **KEGG (May 2023)** | 23,938,003 | 21,910,097 | 11,150,770 | 11,221,753 | 38,627 | 24,784 |
| **InterPro** | 65,155,323 | 78,308,467 | 61,094,216 | 61,446,881 | 102,590 | 25,173 |
| **OrthoDB** | 23,571,268 | 37,141,124 | 22,040,075 | 22,629,213 | 67,414 | 26,865 |
| **EggNOG[a]** | 6,909,072 | 171,141,534 | 6,630,757 | 6,706,098 | 70,417 | 21,067 |
| **PANTHER** | 1,703,797 | 598,660 | 1,351,587 | 1,351,587 | 39,802 | 15,638 |

[a]COG, KOG, and arCOG

**Table S2:** KEGG Orthology average classification performance**.** Best Micro and Weighted F1 for each fraction of unknown is in bold.

| | | Fraction of labeled sequences | | | | |
|---|---|---|---|---|---|---|
| **Metric** | **Model** | **0.05** | **0.25** | **0.5** | **0.75** | **0.95** |
| **F1 Weighted** | **KofamScan** | 0.848 | 0.858 | 0.897 | 0.944 | 0.981 |
| | **FAMUS** | 0.879 | **0.892** | **0.918** | **0.956** | **0.985** |
| | **FAMUS Light** | 0.865 | 0.882 | 0.910 | 0.952 | 0.983 |
| | **HMM Classifier** | **0.927** | 0.884 | 0.838 | 0.794 | 0.764 |
| **F1 Micro** | **KofamScan** | 0.864 | 0.889 | 0.921 | 0.953 | 0.978 |
| | **FAMUS** | 0.888 | **0.909** | **0.930** | **0.959** | **0.980** |
| | **FAMUS Light** | 0.874 | 0.900 | 0.924 | 0.955 | 0.979 |
| | **HMM Classifier** | **0.917** | 0.849 | 0.768 | 0.687 | 0.625 |
| **Precision** | **KofamScan** | **0.981** | **0.976** | **0.967** | **0.932** | **0.738** |
| | **FAMUS** | 0.961 | 0.955 | 0.942 | 0.913 | 0.733 |
| | **FAMUS Light** | 0.959 | 0.955 | 0.941 | 0.909 | 0.726 |
| | **HMM Classifier** | 0.916 | 0.822 | 0.668 | 0.427 | 0.112 |

**Table S3**: InterPro protein families average classification performance. performance for each fraction of unknown for each metric is in bold.

| | | Fraction of labeled sequences | | | | |
|---|---|---|---|---|---|---|
| **Metric** | **Model** | **0.05** | **0.25** | **0.5** | **0.75** | **0.95** |
| **F1 Weighted** | **InterProScan** | **0.904** | **0.868** | 0.834 | 0.808 | 0.797 |
| | **FAMUS** | 0.837 | 0.835 | 0.847 | **0.864** | **0.886** |
| | **FAMUS Light** | 0.838 | 0.832 | **0.849** | 0.861 | 0.883 |
| | **HMM Classifier** | 0.878 | 0.772 | 0.659 | 0.558 | 0.493 |
| **F1 Micro** | **InterProScan** | **0.903** | **0.849** | 0.788 | 0.724 | 0.675 |
| | **FAMUS** | 0.843 | 0.836 | 0.825 | **0.815** | **0.807** |
| | **FAMUS Light** | 0.844 | 0.833 | **0.826** | 0.811 | 0.803 |
| | **HMM Classifier** | 0.875 | 0.754 | 0.608 | 0.461 | 0.344 |
| **Precision** | **InterProScan** | **0.904** | 0.823 | 0.685 | 0.455 | 0.124 |
| | **FAMUS** | 0.902 | **0.850** | **0.751** | **0.559** | 0.186 |
| | **FAMUS Light** | 0.897 | 0.845 | 0.749 | 0.552 | **0.181** |
| | **HMM Classifier** | 0.874 | 0.734 | 0.535 | 0.297 | 0.065 |

**Table S4:** OrthoDB average classification performance**.** performance for each fraction of unknown for each metric is in bold.

| | | Fraction of labeled sequences | | | | |
|---|---|---|---|---|---|---|
| **Metric** | **Model** | **0.05** | **0.25** | **0.5** | **0.75** | **0.95** |
| **F1 Weighted** | **FAMUS** | 0.777 | **0.806** | **0.858** | **0.910** | **0.950** |
| | **FAMUS Light** | 0.769 | 0.797 | 0.848 | 0.908 | **0.950** |
| | **HMM Classifier** | **0.789** | 0/744 | 0.692 | 0.645 | 0.604 |
| **F1 Micro** | **FAMUS** | **0.782** | **0.812** | **0.853** | **0.892** | 0.918 |
| | **FAMUS Light** | 0.775 | 0.803 | 0844 | 0.889 | **0.919** |
| | **HMM Classifier** | 0.778 | 0.704 | 0.611 | 0.521 | 0.442 |
| **Precision** | **FAMUS** | **0.828** | **0.809** | **0.772** | **0.679** | **0.329** |
| | **FAMUS Light** | 0.820 | 0.799 | 0.762 | 0.674 | 0.327 |
| | **HMM Classifier** | 0.775 | 0.670 | 0.508 | 0.296 | 0.066 |

**Table S5:** EggNOG average classification performance**.** performance for each fraction of unknown for each metric is in bold.

| | | Fraction of labeled sequences | | | | |
|---|---|---|---|---|---|---|
| **Metric** | **Model** | **0.05** | **0.25** | **0.5** | **0.75** | **0.95** |
| **F1 Weighted** | **FAMUS** | **0.854** | **0.863** | **0.888** | **0.920** | 0.951 |
| | **FAMUS Light** | 0.776 | 0.789 | 0.840 | 0.899 | **0.952** |
| | **HMM Classifier** | 0.802 | 0.766 | 0.736 | 0.704 | 0.680 |
| **F1 Micro** | **FAMUS** | **0.853** | **0.869** | **0.886** | **0.904** | 0.920 |
| | **FAMUS Light** | 0.768 | 0.799 | 0.844 | 0.887 | **0.926** |
| | **HMM Classifier** | 0.784 | 0.725 | 0.661 | 0.587 | 0.526 |
| **Precision** | **FAMUS** | **0.923** | **0.903** | **0.855** | **0.738** | 0.356 |
| | **FAMUS Light** | 0.881 | 0.852 | 0.816 | 0.705 | **0.361** |
| | **HMM Classifier** | 0.788 | 0.693 | 0.547 | 0.329 | 0.078 |

**Table S6:** KEGG Orthology FAMUS and KofamScan average runtime performance in minutes. The shortest runnning times of each scenario are marked in bold.

| **Sample size** | **5,000** | **10,000** | **15,000** | **20,000** | **25,000** |
|---|---|---|---|---|---|
| **FAMUS CPUs** | 10.20 | 18.19 | 25.11 | 30.13 | 36.02 |
| **FAMUS GPU** | 6.98 | 12.68 | 17.99 | 23.65 | 28.81 |
| **FAMUS Light CPUs** | 7.00 | 11.86 | 16.78 | 19.34 | 23.54 |
| **FAMUS Light GPU** | 4.09 | 7.51 | 10.74 | 14.05 | 17.30 |
| **HMM Classifier** | **4.311** | **7.06** | **10.27** | **13.61** | **16.96** |
| **KofamScan** | 4.56 | 9.44 | 12.67 | 17.54 | 20.72 |

**Table S7:** InterPro FAMUS and InterProScan average runtime performance in minutes. The shortest runnning times of each scenario are marked in bold.

| **Sample size** | **5,000** | **10,000** | **15,000** | **20,000** | **25,000** |
|---|---|---|---|---|---|
| **FAMUS CPUs** | 9.27 | 16.99 | 24.21 | 31.49 | 39.77 |
| **FAMUS GPU** | 7.37 | 13.48 | 18.74 | 25.30 | 31.41 |
| **FAMUS Light CPUs** | 4.74 | 9.05 | 12.73 | 16.23 | 20.50 |
| **FAMUS Light GPU** | 2.85 | 5.26 | 7.19 | 9.92 | 12.27 |
| **HMM Classifier** | **2.56** | **4.93** | **7.01** | **9.54** | **12.02** |
| **InterProScan** | 11.00 | 16.89 | 21.05 | 32.53 | 35.19 |

### Supplementary note 1:

The HMM Classifier used as a benchmark for model performance was comprised of one pHMM per input protein family in the training set. Bit scores were generated for each sequence in the families' respective test sets using hmmsearch from HMMER 3.2.2[1] with default parameters. Each sequence was assigned to the family corresponding to the pHMM with the highest score, if one existed. Otherwise, sequences were labelled as "unknown".
